# Temporal filtering of luminance and chromaticity in macaque visual cortex

**DOI:** 10.1101/2021.03.22.436512

**Authors:** Gregory D. Horwitz

## Abstract

The visibility of a periodic light modulation depends on its temporal frequency and spectral properties. Contrast sensitivity is highest at 8–10 Hz for modulations of luminance but is substantially lower for modulations between equiluminant lights. This difference between luminance and chromatic contrast sensitivity is rooted in retinal filtering, but additional filtering occurs in the cerebral cortex. To measure the cortical contributions to luminance and chromatic temporal contrast sensitivity, signals in the lateral geniculate nucleus (LGN) were compared to the behavioral contrast sensitivity of macaque monkeys. Long wavelength-sensitive (L) and medium wavelength-sensitive (M) cones were modulated in phase, to produce a luminance modulation (L+M), or in counterphase, to produce a chromatic modulation (L-M). The sensitivity of LGN neurons was well matched to behavioral sensitivity at low temporal frequencies but was approximately 7 times greater at high temporal frequencies. Similar results were obtained for L+M and L-M modulations. These results show that differences in the shapes of the luminance and chromatic temporal contrast sensitivity functions are due almost entirely to pre-cortical mechanisms. Simulations of cone photoreceptor currents show that temporal information loss in the retina and at the retinogeniculate synapse exceeds cortical information loss under most of the conditions tested.

## Introduction

Signal processing in the visual system preserves of some types of information while eliminating others. If perfect knowledge of neuronal activity at one stage of the visual system (e.g. visual cortex) allows for perfect reconstruction of activity at an earlier stage (e.g. the photoreceptors), then information is perfectly preserved between them. If, instead, multiple patterns of activity at an early stage produce indistinguishable patterns at a later stage, then information has been lost. The ability of an observer to detect a stimulus—to distinguish it from a blank—reflects information loss accumulated throughout the visual system. A central goal of visual neuroscience is to understand which types of information are lost at which stage of the visual system to mediate stimulus detection.

A salient example of information loss in the visual system is evident in the fact that the visibility of a periodic stimulus depends on its temporal frequency. This relationship, the temporal contrast sensitivity function, plays important roles in industry (Legall 1991) and medicine (Owsley 2011, Tyler 1981), but its biological basis is incompletely understood. This uncertainty is due in part to methodological differences between neurophysiological and behavioral studies. Temporal contrast sensitivity depends on many factors including: observer species, background luminance, retinal eccentricity, stimulus size, and duration (Benardete & Kaplan 1997b, Benardete & Kaplan 1999a, Lee et al. 1990, Lindbloom-Brown et al. 2014, Merigan 1980, Pokorny et al. 2001, Snowden & Hess 1992, Snowden et al. 1995, Solomon et al. 2002, Swanson et al. 1987, Van der Horst 1969). In the current study, care was taken to match these factors between neurophysiological and behavioral measurements, providing a clearer picture of their relationship than has previously been available.

Chromatic modulations are easier to see than luminance modulations at low temporal frequencies, but at higher frequencies, the reverse is true (De Lange Dzn 1958, Kelly & van Norren 1977)(Figure 1A). The photoreceptors cannot be responsible for this difference because the same photoreceptor types that underlie luminance detection also underlie chromatic detection. Luminance stimuli modulate the long- (L) and medium wavelength-sensitive (M) cones in phase (L+M), whereas most chromatic stimuli modulate them in counterphase (L-M). Differences in the temporal filtering of these two stimulus classes must therefore be due to stages of the visual system where signals from the L- and M-cones are already mixed.

**Figure 1.**
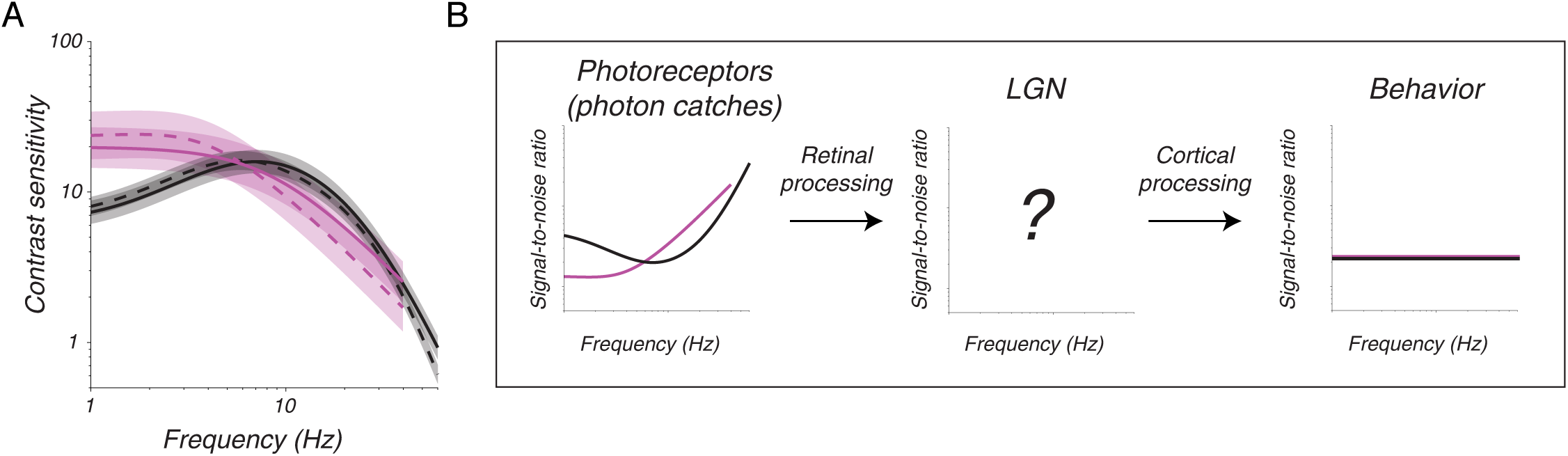
**A:** Temporal contrast sensitivity functions from monkey 1 (dashed) and monkey 2 (solid) for L+M modulations (black) and L-M modulations (magenta). Curves represent the means across receptive field locations of recorded LGN neurons, and bands represent ± 1 standard deviation. **B:** Schematic of the experimental logic. A set of stimuli that varied in relative L- and M-cone phase (L+M or L-M) and temporal frequency was presented at the RF of each neuron studied. The contrast of each stimulus was adjusted (left) so that the signal-to-noise ratio at the level of behavior (right) was fixed. Predictions in the middle panel are fuzzy to depict uncertainty in the signal-to-noise ratio of responses in the LGN.

The purpose of this study was to quantify cortical and pre-cortical contributions to luminance and chromatic temporal contrast sensitivity. Cortical contributions were computed by comparing the behavioral sensitivity of a macaque monkey to that of a computational observer of spikes in the lateral geniculate nucleus (LGN)(Figure 1B). Pre-cortical contributions were computed by comparing two computational observers: one of LGN spikes, and one of simulated currents across the outer segments of modelled cone photoreceptors.

The main result of these comparisons was that information loss in the cortex was similar for L+M and L-M modulations, whereas information loss between the cones and LGN differed profoundly for L+M and L-M modulations. Differences in luminance and chromatic temporal contrast sensitivity are therefore due to processes occurring upstream of the LGN with minimal cortical involvement.

## Results

Two monkeys (M. mulatta) performed a 2-alternative, forced-choice contrast detection task that required them to report on which side of a computer screen a drifting Gabor stimulus appeared. Detection thresholds were measured as a function of stimulus location, temporal frequency, and the amplitude of L- and M-cone modulations. A model was developed that predicted detection thresholds as a function of all of these parameters jointly (Gelfand & Horwitz 2018 and Figure 1A). Visual stimuli were constructed on the basis of this model and used to measure the signal-to-noise ratio (SNR) of LGN neuronal responses (Figure 1B). All stimuli were at the monkeys’ behavioral detection threshold, or equivalently, matched for SNR at the output of the visual system.

### LGN single-unit responses

The spatial and spectral sensitivity of each recorded LGN neuron were characterized with a white-noise stimulus (Horwitz 2020). Spike-triggered averaging was used to locate the receptive field (RF) center and to identify the physiological type of each neuron. Fifteen neurons were classified as magnocellular (8 from monkey 1 and 7 from monkey 2) and 38 as parvocellular (19 from each monkey). Each recorded neuron was then stimulated with Gabor patterns centered on the RF that varied across trials in temporal frequency and L- and M-cone modulation phase (in-phase, L+M, or counterphase, L-M). The L- and M-cone contrasts were always equal, and their maximum was set by the limits of the display (0.19 for the L-M stimulus and 0.86 for the L+M stimulus). A blank stimulus was included to measure baseline firing statistics.

A representative magnocellular neuron responded to L+M modulations vigorously at high temporal frequencies and more weakly as temporal frequency was reduced (Figure 2A) (for similar data from a second magnocellular neuron see Horwitz 2020). This neuron also responded to L-M modulations but only at the highest frequencies tested and then only transiently (Figure 2B). An example parvocellular neuron responded more vigorously to L-M modulations than to L+M modulations (Figures 2C and 2D), although, as expected from their low contrast, none of the stimuli used in this study drove parvocellular neurons strongly (the example in Figure 2C & 2D is among the most responsive parvocellular neurons in the data set).

**Figure 2.**
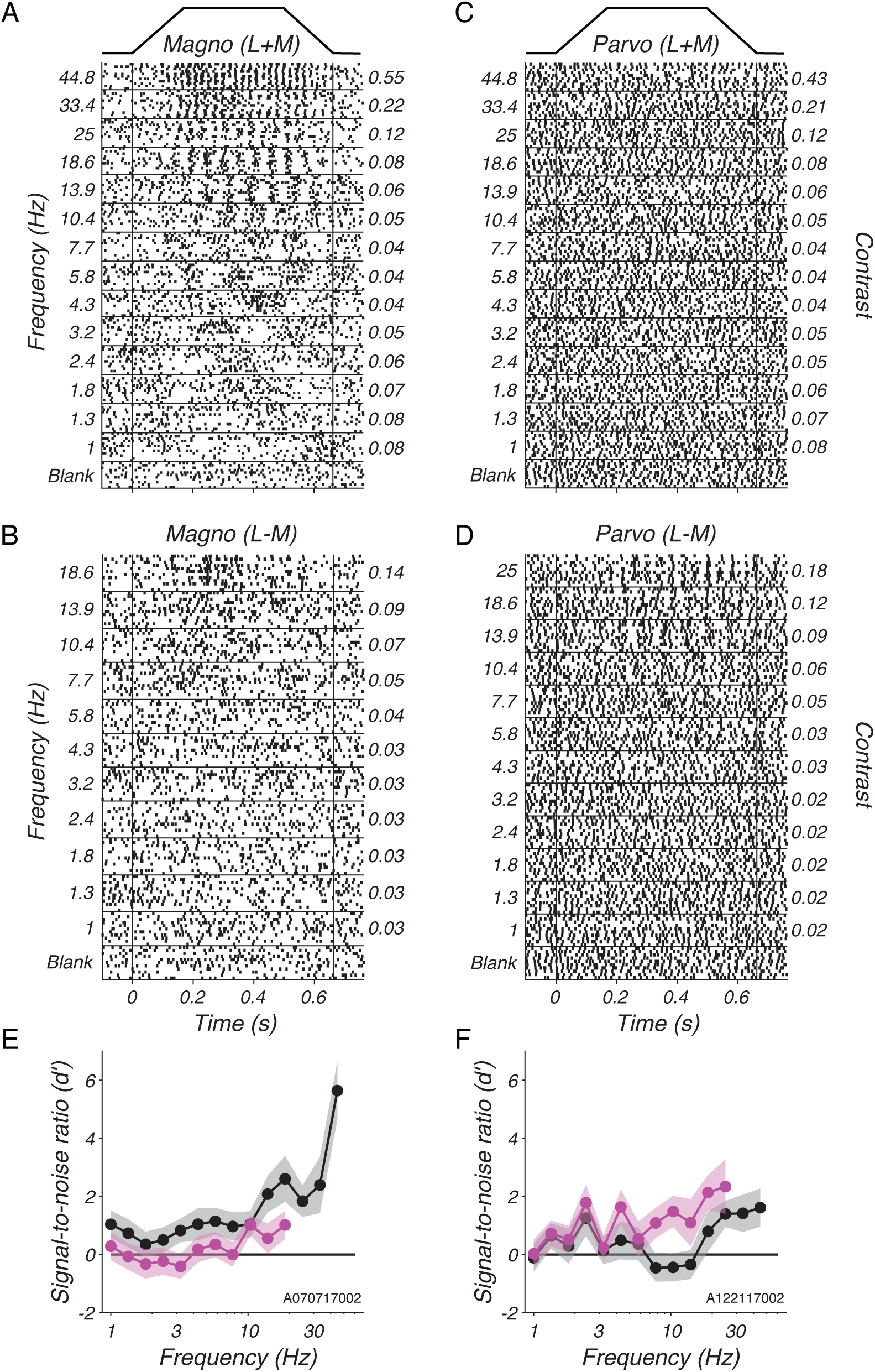
Responses of two LGN neurons to Gabor stimuli near behavioral detection threshold. **A**: Raster plot of magnocellular responses to L+M modulations. Trials have been sorted by temporal frequency (left ordinate) which covaries with cone contrast (right ordinate, identical for L- and M-cones) to maintain a constant level of stimulus detectability. The temporal envelope of the Gabor stimulus is shown above the rasters. **B**: Identical to A but showing responses to L-M modulations. **C & D**: Identical to A & B but for a parvocellular neuron. **E**: Signal-to-noise ratio (d’) calculated from responses in panel A (black) and from responses in B (magenta). Points represent means, and shaded bands represent ±1 standard error estimated by non-parametric bootstrap. **F**: Identical to E but for the parvocellular neuron.

The SNR of each response was calculated by comparing it to baseline activity. This analysis assumes that the signal in the spike trains is at the fundamental temporal frequency of the stimulus (see Methods), but this assumption was not critical to the main results (see Supplemental Information, Supplemental Figures 1 & 2). The example magnocellular neuron had greater SNR for L+M than for L-M modulations at all frequencies tested (Figure 2E). The example parvocellular neuron had greater SNR for L-M modulations than for L+M modulations above 6 Hz (Figure 2F).

The relationships among spiking responses, temporal frequency, L- and M-cone modulation, and cell type become clearer when data are averaged across neurons (Figure 3). As expected, magnocellular neurons were more sensitive to L+M modulations, and parvocellular neurons were more sensitive to L-M modulations (Wiesel & Hubel 1966). The SNR of magnocellular and parvocellular responses increased smoothly from 1 to 20 Hz despite the fact that contrast changed with in temporal frequency in different ways for L+M and L-M modulations over this range to keep each stimulus at detection threshold.

**Figure 3.**
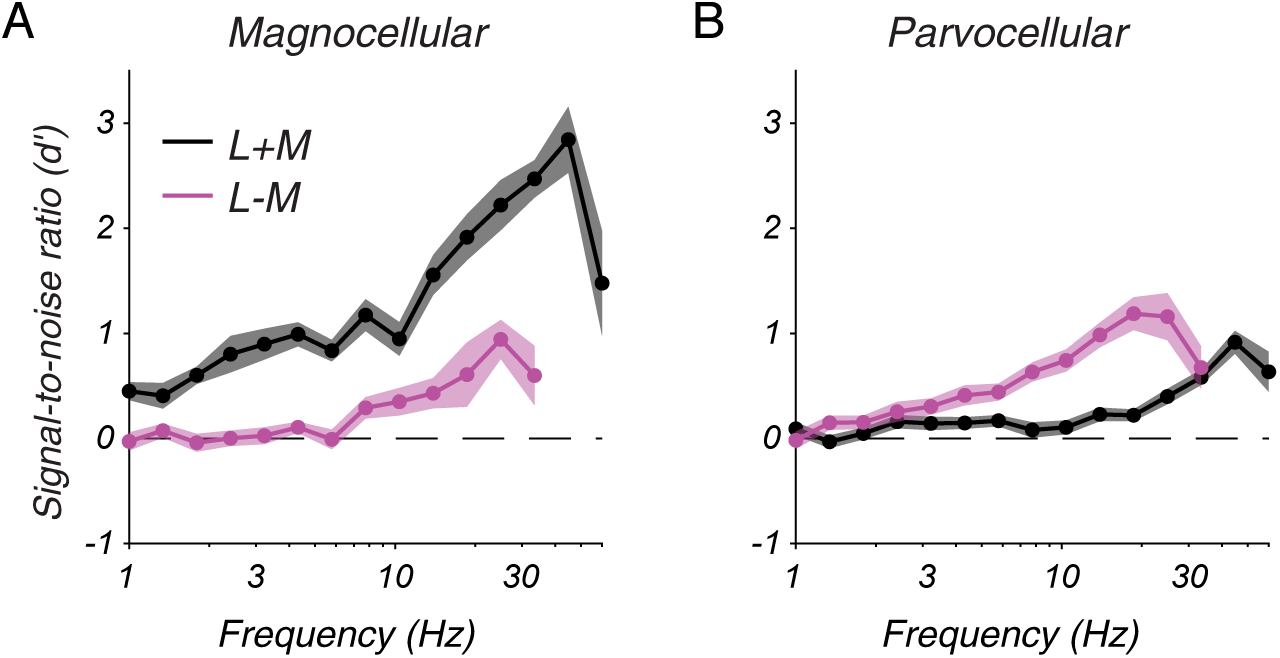
**A**: Signal-to-noise ratio (d’) averaged across magnocellular neurons (A) and parvocellular neurons (B). Shaded bands represent ±1 standard error of the mean.

The SNR of the average neuron (Figure 3) is lower than the SNR of neuronal populations. To estimate the SNR of a population of LGN neurons, the SNR of individual LGN neurons was inflated by an estimate of how many LGN neurons were modulated by the stimulus, as described in the next section.

### Population SNR analysis

Magnocellular neurons have greater contrast sensitivity than parvocellular neurons do at matched eccentricity, but they are less numerous, raising the possibility that, as populations, parvocellular neurons might have greater SNR (Croner & Kaplan 1995). To estimate the SNR of neuronal populations, a model was constructed using parameters taken from the literature, without fitting to data (Horwitz 2020). The model provided a scale factor for each neuron that reflects how many times greater the SNR of a population of similarly sensitive neurons is expected to be. Scale factors were 2.1-fold greater (±0.4 SD) for parvocellular neurons than magnocellular neurons at matched eccentricity.

Parvocellular population SNR rose steeply with the temporal frequency of L-M modulations, and magnocellular population SNR rose similarly with the temporal frequency of L+M modulations (Figure 4A). Magnocellular and parvocellular populations were also weakly and similarly responsive to their non-preferred modulations, L-M and L+M, respectively. The similarity of these patterns is striking considering that these data were derived from recordings from two distinct populations of neurons responding to two sets of stimuli that varied in temporal frequency and L- and M-cone contrast in different ways.

**Figure 4.**
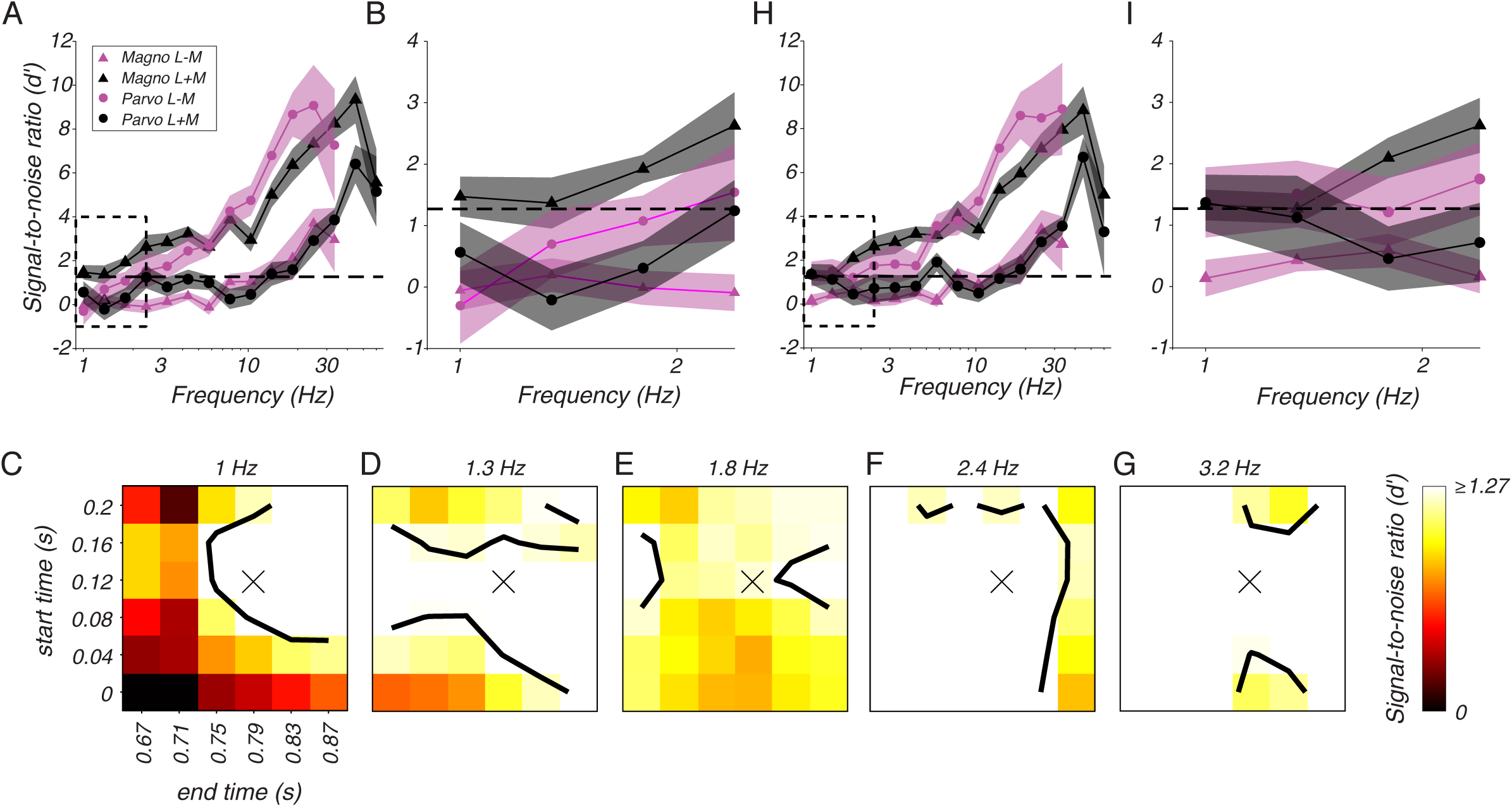
Population signal-to-noise analysis. **A**: Population signal-to-noise ratio (d’) as a function of temporal frequency for magnocellular neurons (triangles) and parvocellular neurons (circles) in response to L+M modulations (black) and L-M modulations (magenta). Bands represent ±1 standard error of the mean across neurons. Dashed line at 1.27 indicates the d’ at the level of perceptual decision-making inferred from behavioral sensitivity. Dashed rectangle represents region magnified in **B**. **C**: Population d’ for parvocellular neurons in response to 1 Hz, L-M modulations as a function of the start time (ordinate) and end time (abcsissa) of the spike counting window. Contour is drawn at d’ = 1.27. A spike counting window delayed by 120 ms from the stimulus presentation epoch (marked by an "X") produced a greater d’ value than the window used in A & B, which did not take response latency into account (lower left corner). **D– G**: identical to C but for 1.3, 1.8, 2.4, and 3.2 Hz modulations, respectively. **H & I**: identical to A & B but counting spikes from 120 ms after the stimulus appeared until 120 ms after the stimulus disappeared.

To quantify how much information was lost in the cortex, SNR in the LGN was compared to behavioral sensitivity. For this purpose, the monkeys’ performance at threshold, 82% correct, was converted to an SNR of 1.27 (Green & Swets 1966, see Methods). At high frequencies, the SNR of magnocellular and parvocellular neurons exceeded this level by approximately 7-fold in response to L+M and L-M modulations, respectively. At lower frequencies, SNR in the LGN was lower. In fact, at the lowest frequencies tested, parvocellular population SNR fell below behavioral SNR for both L+M and L-M modulations (Figure 4B and Methods). Parvocellular neurons are the sole conduit by which low temporal-frequency L-M modulations are transmitted from the eye to the cortex, so parvocellular SNR was underestimated.

The analysis in Figure 4A & 4B was based on spikes recorded between stimulus onset and disappearance, including the slow (166 ms) contrast ramps at the beginning and end of each stimulus presentation. No adjustment was made for response latency, which biased SNR downward. To examine the effects of spike counting window on SNR, the start and stop times for spike inclusion were varied independently over a 200-ms range (Figure 4C–4G). This analysis showed that delaying the spike counting window relative to the stimulus presentation by ∼120 ms boosted parvocellular population SNR sufficiently to mediate behavior at even the lowest temporal frequencies tested. This delay presumably reflects the low contrast sensitivity of parvocellular neurons combined with the slow contrast increase at the beginning of each stimulus presentation.

Across cell types and stimulus conditions, delaying the spike counting window by 120 ms affected SNR only subtly (compare Figure 4A to 4H and Figure 4B to 4I). Over a broader range of spike counting windows, none was found that rendered parvocellular populations significantly more sensitive to low-frequency L-M modulations than the monkey (Supplementary Figure 3). Over the same range of windows, magnocellular and parvocellular population SNR were similar for L+M and L-M modulations, respectively (Supplementary Figure 3).

Three conclusions can be drawn from these analyses: low-frequency information is preserved with near-perfect fidelity downstream of the LGN, the amount of information loss downstream of the LGN changes smoothly with temporal frequency, and the amount of information lost downstream of the LGN is nearly independent of whether L- and M-cone modulations are in-phase or counterphase. The difference between the luminance and chromatic temporal contrast sensitivity functions is therefore due primarily to information loss upstream of the LGN, which is quantified next.

### SNR loss upstream of the LGN

To measure how much information was lost between the cone photoreceptors and the LGN, cone photocurrent responses to the stimuli used in the LGN recordings were simulated using the model of Angueyra and Rieke (2013). SNR loss between the cones and the LGN exceeded SNR loss in the cortex and was particularly severe at low temporal frequencies (Figure 5, diagonal cross hatches). Only 5% of the SNR available in cone outer segment currents in response to low-frequency L+M modulations reached the LGN (Figures 5A & 5B). In response to L-M modulations, information transmission efficiency was more than doubled (Figures 5C & 5D) (see also Chaparro et al. 1993). Above 5 Hz, the situation reversed; SNR loss for L-M modulations exceeded SNR loss for L+M modulations. This analysis confirms differential retinal filtering of L+M and L-M modulations and shows that most of the information loss under the conditions tested occurred upstream of the LGN.

**Figure 5.**
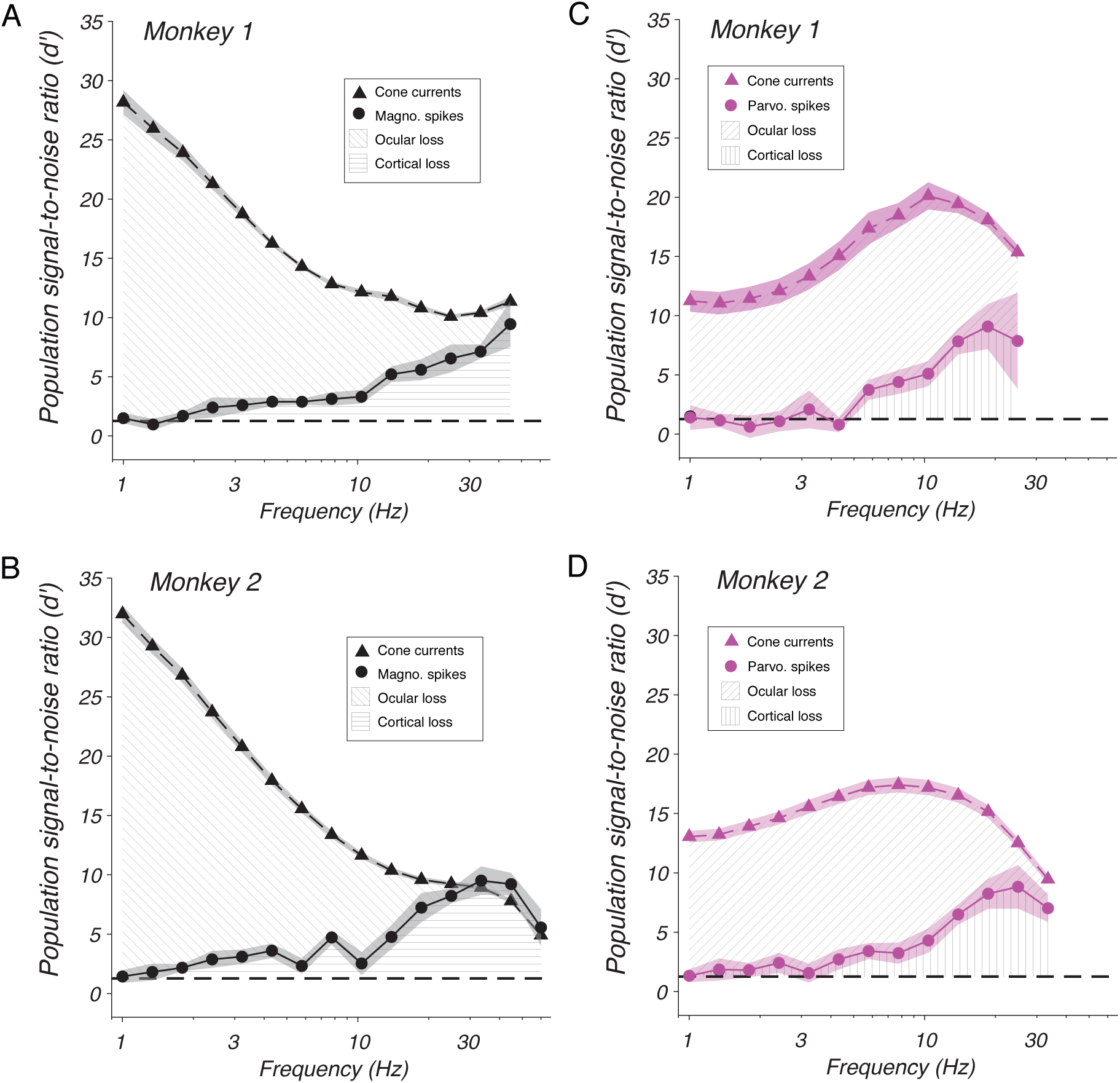
Population signal-to-noise ratio (d’) for monkey 1 (**A & C**) and monkey 2 (**B & D**). Symbols represent means across neurons, and shaded bands represent ± 1 standard error of the mean. Population d’ was calculated from simulated cone currents (triangles) and recorded LGN spikes (circles) in response to L+M modulations (black) and L-M modulations (magenta). Diagonal cross-hatching shows the difference in d’ between cone currents and LGN spikes. Horizontal and vertical cross-hatching shows the difference in d’ between LGN spikes and behavior.

## Summary and Discussion

Much of the information in the light absorbed by photoreceptors fails to reach perception (Barlow 1957, Geisler 1989, Geisler 2011). Identifying where and how this information is lost is a key step towards understanding the biological basis of vision. The distinctive temporal properties of luminance and chromatic vision offer insight into this broader issue. The fact that information loss is temporal, not spatial, indicates a neural basis as opposed to an optical one. The fact that the same photoreceptor types mediate both aspects of vision indicates that the information loss is downstream of the photoreceptors. Previous studies have shown that low-frequency L+M modulations are selectively filtered in the retina, and that high-frequency modulations are filtered in the cortex (Kaplan & Benardete 2001, Kaplan et al. 1990). The new contributions of the current study are the quantitative comparison of information loss upstream and downstream of the LGN and the demonstration that cortical filtering of L+M and L-M modulations is similar across temporal frequencies.

### Mechanisms of SNR loss in the retina and LGN

The stimuli used in this study had little spatial structure and were approximately uniform within the RF of each LGN neuron studied. Consequently, center-surround antagonism reduced SNR in response to L+M modulations at low temporal frequencies (Figure 6A & 6B, top). At higher temporal frequencies, the delay of the surround became an appreciable fraction of the stimulus period, causing excitation from the center to move closer in time to the release of surround inhibition (Enroth-Cugell et al. 1983, Robson 1966)(Figure 6B, bottom). This change in the relative timing of excitation and inhibition largely explains the weaker response of LGN neurons to low frequency L+M modulations than to higher frequency (5–10 Hz) L+M modulations (Benardete & Kaplan 1997a, Benardete & Kaplan 1999a) (Figure 6C).

**Figure 6.**
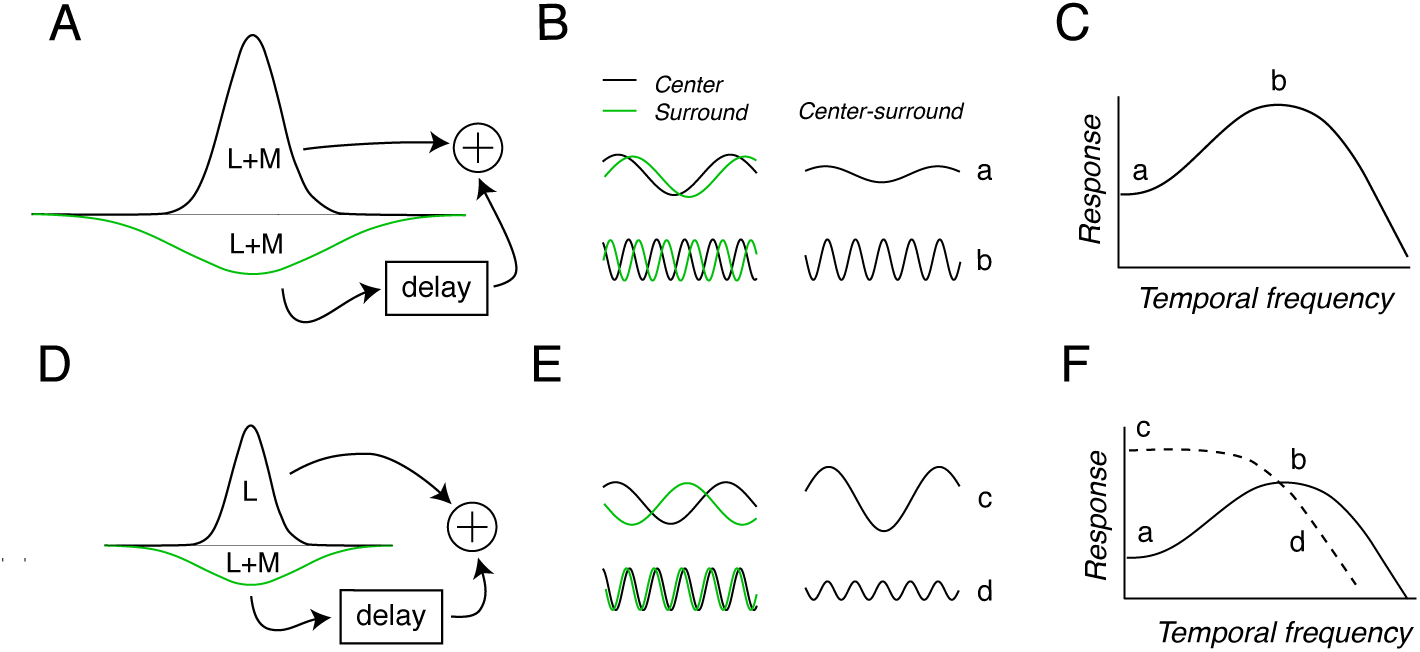
Temporal filtering by center-surround receptive field antagonism. **A**: Schematic receptive field profile of an ON-center cell. Center (narrow upright Gaussian) and surround (broad upside-down Gaussian) are sensitive to a sum of L- and M-cone modulations. **B**: Modulations of the center (black) and surround (green) in responses to L+M modulations (left) are subtracted (right) to represent the net response to a stimulus that modulates both center and surround together. **C:** Temporal frequency tuning of the neuron in **A**. **D–F:** Similar to A–C but for an L-ON cell. Traces in E represent responses to L-M modulations. Dashed curve in F represents temporal frequency tuning for L-M modulations. a, b, c, and d in B & E denote stimuli that correspond to points on the temporal frequency tuning curves in C & F.

Most parvocellular neurons with parafoveal RFs receive input from a single cone type to the center of their RFs and a mixture of L- and M-cones to the surround. For these neurons, L-M modulations invert the influence of the surround relative to the center. A parvocellular L-ON neuron, for example, is excited by an increase in L-cone contrast at the center and disinhibited by a decrease in M-cone contrast in the surround (Figure 6D). When close together in time, these influences combine to drive a strong response (Figure 6E, top). When the temporal frequency of the modulation is sufficiently high that excitation from the center and inhibition from the surround coincide, the response is reduced (Figure 6E, bottom & 6F). This explains the low-pass temporal frequency tuning of parvocellular neurons to L-M modulations (Benardete & Kaplan 1999b, Lankheet et al. 1998).

The high-frequency roll-off of magnocellular and parvocellular responses is due largely to phototransduction, the dynamics of which depend on mean light intensity. Across a broad range of light levels, increasing the mean intensity of a modulated light increases the speed of cone responses (Baudin et al. 2019), retinal ganglion cell (RGC) responses (Purpura et al. 1990), and shifts the peak of the psychophysical temporal contrast sensitivity function to higher frequencies (De Lange Dzn 1958). The ability to predict the shape of the high-frequency limb of the temporal contrast sensitivity function on the basis of the cone temporal impulse response across light levels suggests that that cortical filtering is independent of light level (Lee et al. 1990, Rider et al. 2019, Stockman et al. 2006).

Temporal filtering at the retinogeniculate synapse appears to be modest under most conditions (Alitto & Usrey 2008, Benardete & Kaplan 1997a, Benardete & Kaplan 1999b, Kaplan et al. 1987, Kaplan & Shapley 1986). Many of the stimuli used in the current study had low contrast, making retinogeniculate transmission particularly efficient (Kaplan et al. 1987). High-frequency stimuli had higher contrasts, but the similarity in SNR of cone currents and LGN neurons at these frequencies suggests that information loss at the retinogeniculate synapse was minimal.

### Mechanisms of SNR loss in the cortex

The SNR gap between the LGN and behavior is due, at least in part, to processes occurring in area V1 (Hawken et al. 1996). One mechanism that may contribute to high-frequency filtering in V1 is push-pull excitation-inhibition (Tolhurst & Dean 1990). Simple cells in V1 receive spatially coincident excitation and inhibition that prevent high-contrast, non-preferred stimuli from driving a response (Troyer et al. 1998) and reduce sensitivity to high temporal frequency modulations (Krukowski & Miller 2001, Krukowski et al. 2001). An intuition for the latter effect is that excitation and inhibition cancel when triggered simultaneously. The dominant inhibition required by the push-pull model ensures that cancellation is complete. The slow kinetics of NMDA-sensitive channels in V1 neurons broaden the window of effective simultaneity (Eickhoff et al. 2007, Lester et al. 1990).

Most of the data supporting the push-pull model are from cat, but the same principles are likely at work in primates (Conway & Livingstone 2006, Kremkow & Alonso 2018). Monkeys have luminance-tuned simple cells, like cats do, but unlike cats, monkeys have a large population of cone-opponent V1 neurons. Some of these cone-opponent neurons combine visual signals antagonistically and roughly linearly across their RFs, consistent with the push-pull model (Conway & Livingstone 2006, De & Horwitz 2021). One possibility that is consistent with the results of this study is that push-pull excitation-inhibition reduces the SNR of high-frequency cone-opponent and non-opponent modulations similarly in V1.

Some V1 neurons respond to high-frequency signals that cannot be detected, implying a high-frequency filter within or downstream of V1 (Engel et al. 1997, Gur & Snodderly 1997, Hawken et al. 1996, Jiang et al. 2007, Krolak-Salmon et al. 2003, Shady et al. 2004, Vul & MacLeod 2006, Williams et al. 2004, Zhigalov et al. 2019). The possibility therefore remains that high-frequency modulations are conducted from the output of V1 to decision-making circuitry efficiently. Laminar V1 recordings could be used to test this hypothesis.

### Relationship to previous work

Two innovations set the current study apart from those previous. The first was holding fixed several factors between neurophysiological and behavioral measurements: the species and identities of the subjects, the intensity of the display background, the retinal eccentricity of the stimulus, and the stimulus size. Two previous primate studies matched these parameters, but neither of them varied temporal frequency, and the one that varied color reported data from few neurons (Jiang et al. 2015a, Jiang et al. 2015b, Sperling et al. 1978). A second innovation was the use a cone current model to quantify information loss through the retina and retinogeniculate synapse (Angueyra & Rieke 2013, Hass et al. 2015, Horwitz 2020).

Results from this study are broadly consistent with those of Lee et al. (1990) who compared contrast detection thresholds of human observers to the responses of individual magnocellular-projecting (M) and parvocellular-projecting (P) RGCs. M RGCs responded strongly to luminance modulations and weakly to chromatic modulations. The reverse was true for P RGCs. Individual RGCs of both types were less sensitive than human observers at low frequencies and more sensitive at high frequencies. Results of the current study extend these observations by showing that the sensitivity of LGN populations and observers match at low temporal frequencies, that the SNR of M and P populations are similar across temporal frequencies at contrast detection threshold, and that retinal circuitry is lossier than cortical circuitry except at high frequencies.

The idea that L-M and L+M temporal contrast sensitivity functions can be directly related to the activity in the M and P pathways has been the subject of much debate. Single unit recordings are ill-suited for settling this debate because, as shown in this study, many stimuli activate both pathways even at detection threshold. The only stimulus that achieved decisive pathway-specificity in this study was the low temporal frequency, L-M stimulus, which modulated parvocellular neurons weakly but exclusively. Low-frequency L+M stimuli modulated magnocellular neurons more strongly than parvocellular neurons, but both populations carried measureable signal. At high frequencies, both magnocellular and parvocellular neurons responded briskly to L+M and L-M stimuli. In lesioned animals, high temporal frequency modulations are detected via the magnocellular pathway, at least at low spatial frequencies (Merigan & Eskin 1986, Merigan & Maunsell 1990, Schiller et al. 1990).

### Spatial contrast sensitivity

Visual sensitivity under a range of conditions is bandpass for luminance contrast and low pass for chromatic contrast. Interestingly, this pattern is consistent whether modulations are temporal or spatial. A normative explanation is that L-M signals in natural scenes are small (Ruderman et al. 1998) but important (Carvalho et al. 2017, Rosenthal et al. 2018). Detecting these signals is facilitated by integration (low pass filtering), a strategy that works over space or time due to the large, stationary nature of objects. L+M signals in natural scenes have greater amplitude, so they can be detected with less integration, permitting finer spatial and temporal resolution and the consequent benefits for visually guided action.

Some mechanisms underlying spatial and temporal visual filtering are shared. For example, low-frequency spatial and temporal modulations are filtered via center-surround RF antagonism (Robson 1966), and high-frequency modulations are filtered via phototransduction (Cottaris et al. 2020). The spatial effects of phototransduction are linked to small eye movements produced during fixation. A small displacement of a high spatial-frequency grating can stimulate individual cone photoreceptors with contrast increments and decrements close together in time, causing cancellation.

Other mechanisms of spatial and temporal filtering differ, one of which is highlighted by the current results. This study showed that temporal filtering of luminance and chromatic modulations is similar in the cortex. In contrast, spatial filtering of luminance and chromatic modulations differs substantially. High spatial-frequency luminance sensitivity is limited by midget ganglion cell density, implying near-perfect fidelity of cortical information transmission (Anderson et al. 1991, Banks et al. 1987, Banks et al. 1991, Dacey 1993). Chromatic spatial sensitivity, on the other hand, is subject to substantial additional filtering in the cortex (Martin et al. 2001, Mullen & Kingdom 2002, Mullen et al. 2005, Solomon et al. 2005).

### Caveats

Several disparate data sets were converted to a common SNR metric to facilitate comparison across stages of the visual system. This conversion required mathematical models that could lead to erroneous conclusions if based on erroneous assumptions. The basis of each model, the approximations and assumptions made in their construction, and probable sources of error are discussed below.

### The cone current model

The cone current model was based on patch clamp recordings from *ex vivo* macaque cones under light levels similar to those used in the current study (Angueyra & Rieke 2013). The model approximates current noise as being independent of the signal, which is reasonable at the moderate light levels used in this study (Figure 1 of Angueyra & Rieke 2013). Cone signaling dynamics were approximated as independent of eccentricity, which is reasonable over the range investigated in this study (2–14°) (Sinha et al. 2017). Absolute detection thresholds predicted by this model are close to those measured behaviorally (Angueyra & Rieke 2013, Koenig & Hofer 2011).

Weaknesses of the model include the fact that it is based on a single, canonical temporal impulse response, noise spectrum, and cone density, all of which presumably vary across observers (density does; see Curcio et al. 1987). Indeed, results of this study provide indirect evidence for individual differences. LGN neurons in monkey 2 were more sensitive than those in monkey 1, relative to the cone model (Figure 5). This this was true for both magnocellular and parvocellular neurons, consistent with a systematic underestimate of cone sensitivity in monkey 2.

At the highest frequencies tested, magnocellular SNR slightly exceeded the SNR of simulated cone currents in monkey 2 (Figure 5B). This is unrealistic; SNR cannot increase between the cones and the LGN. The population model is not responsible for this discrepancy. The SNR of individual magnocellular neurons from monkey 2 exceeded the SNR of the simulated cones inside their RFs (Supplemental Figure 4). One explanation is that the number of cones in monkey 2 was underestimated (Packer et al. 1989). Alternatively, the high-frequency sensitivity of the simulated cones may have been underestimated due to the *ex vivo* preparation or the fact that cone current simulations were based on recordings made at 4,000- 6,500 photoisomerizations per second whereas cones in the monkeys’ eyes absorbed approximately 7,400–8,800 photoisomerizations per second during the LGN recording experiments.

### The LGN model

The LGN population model included correlations between neurons of a common type (magnocellular or parvocellular) but not between populations. Consequently, population SNR was computed for magnocellular and parvocellular populations separately. SNR could not be computed for both populations jointly without additional assumptions that are ill-constrained by data.

The SNR of each LGN population is a lower bound on the SNR of both of them together. Note that this lower bound approached the theoretical *upper* bound imposed by SNR of cone outer segments at high temporal frequencies (Figure 5). This leads to a prediction: the signals carried by magnocellular and parvocellular neurons with overlapping RFs are largely redundant in response to high temporal-frequency modulations. This conclusion is consistent with the idea that the L-M signals carried by magnocellular neurons derive from the same circuits that mediate cone-opponency in midget RGCs (Lee & Sun 2009, Stockman et al. 2018). It is also consistent with the fact that the responses of midget and parasol RGCs with overlapping RFs share noise that is inherited from the photoreceptors (Ala-Laurila et al. 2011).

### The behavioral model

The behavioral model was based on 13,760 detection trials from monkey 1 and 28,960 from monkey 2. Contrast sensitivity functions predicted from the model (Figure 1A) were similar to those from the literature and to those measured from human subjects performing the same task in the same testing apparatus (Gelfand & Horwitz 2018, Merigan 1980, Stavros & Kiorpes 2008). The model accurately predicted contrast detection thresholds collected after the electrophysiological experiments (Horwitz 2020). Probable error in estimated behavioral SNR was approximately 30% (see Methods).

### Conclusion

By comparing signal loss upstream and downstream of the LGN quantitatively and under identical conditions, this study showed that the differences between the luminance and chromatic temporal contrast sensitivity functions are due to processes upstream of the LGN with little if any cortical involvement.

## Acknowledgements

I am deeply grateful to Fred Rieke and Juan Angueyra for valuable discussions, EJ Chichilnisky for helpful feedback on the manuscript, Emily Gelfand and Lisa McConnell for excellent animal husbandry and training, and Zack Lindbloom-Brown for computer programming. This study was supported by NIH grants EY018849 and OD010425.

## STAR Methods Resource Availability

Data and code are available at https://github.com/horwitzlab/LGN-temporal-contrast-sensitivity. Requests for additional information not available in that repository will be fulfilled by Lead Contact Greg Horwitz.

## Experimental Model and Subject Details

Two macaque monkeys (M. mulatta, both male) were used in these experiments. Monkeys 1 and 2 were 13 and 7 years old, respectively, at the time of data collection. All procedures were approved by the University of Washington Institutional Animal Care and Use Committee. Monkeys 1 and 2 in the current study are monkeys 1 and 2 from (Gelfand & Horwitz 2018) and from (Horwitz 2020). The data analyzed in this report were collected contemporaneously with the data reported in (Horwitz 2020) and partially overlap with this previous data set.

## Method Details

### Behavioral task

Behavioral detection thresholds were measured using a two-alternative, forced-choice contrast detection task (Gelfand & Horwitz 2018). Monkeys sat in a dark room, 61 cm away from a rear-projection screen that was illuminated by a digital light projector (Propixx, VPixx, Inc.) updating at 240 Hz. Stimuli were generated using routines from the Psychophysics Toolbox (Brainard 1997, Kleiner et al. 2007, Pelli 1997). The spectral power distribution of each display primary was characterized with a PR650 spectroradiometer (PhotoResearch, Inc.). The background was an equal-energy white metamer at 130 cd/m^2^, producing approximate isomerizations per cone per second of L: 8900, M: 7400, and S: 2300.

The monkey initiated each trial by fixating a 0.2 x 0.2° spot at the center of the screen. An upward-drifting, horizontally oriented Gabor stimulus (1 cycle/° in a 0.15° standard deviation envelope) appeared in either the right or left hemifield. Stimulus contrast increased linearly over 166 ms, remained constant for 334 ms, and then ramped down over 166 ms. After a 100–600 ms delay, the fixation point vanished, and saccade targets appeared 2° to the right and left of fixation. The monkey received a liquid reward for making a saccade to the target in the hemifield in which the Gabor stimulus had appeared. Stimulus contrast, color direction in the LM plane, and temporal frequency varied across randomly interleaved trials, and stimulus location varied across days.

The duration of the stimuli used in this study exceeded psychophysical integration times, which are color and temporal frequency-dependent (King-Smith & Carden 1976, Rovamo et al. 2003, Rovamo et al. 1996, Smith et al. 1984). A protracted stimulus was necessary to probe low temporal frequencies; low-frequency stimuli cannot be brief.

The data were fit with a model that predicts detection threshold as a function of temporal frequency, color direction in the LM plane, and location in the visual field. This model was used to construct stimuli that were adjusted in contrast to be near the monkeys’ detection threshold (Gelfand & Horwitz 2018).

### Electrophysiology

Responses of single LGN units were measured with extracellular tungsten electrodes (Fredrick Haer, Inc.) and recorded with a Multichannel Acquisition Processor system (Plexon, Inc.). Spike isolation was performed online with SortClient software and refined offline with OfflineSorter software. Visual fixation was tracked with a scleral search coil (Riverbend Instruments, Inc.) and was required to remain in a 1 x 1° window. Liquid rewards were given for successful fixation. The visual display used in the electrophysiological experiments was identical to that used in the behavioral experiments.

### White-noise stimulation

Each recorded LGN neuron was characterized with a white-noise stimulus that consisted of a 10 x 10 grid of 0.2° pixels. The light at each of these pixels was determined by independent, random draws from red, green, and blue Gaussian intensity distributions. The stimulus updated at 60 Hz (every four frames). Spike-triggered averaging was performed online to locate the receptive field of each recorded neuron and offline to classify it as magnocellular or parvocellular.

### Near-threshold Gabor stimulation of LGN neurons

Following white noise characterization, each neuron was stimulated with a sequence of Gabor patterns centered on its receptive field. All stimuli were equated for detectability using the model of Gelfand and Horwitz (2018). Each neuron had a unique receptive field location and was therefore probed with a unique set of contrasts. Every stimulus modulated the L- and M-cones of the Stockman, MacLeod, and Johnson 10° standard observer with identical contrasts and did not modulate the S-cones. In randomly interleaved trials, L- and M-cone modulations were in phase, to create an L+M stimulus, and in counterphase, to create an L-M stimulus.

## Quantification and Statistical Analysis

### LGN SNR calculation

Firing rate modulations of LGN neurons in response to the stimulus were nearly symmetric around the baseline rate. To quantify the neural response, the modulation amplitude of LGN spike trains at the fundamental frequency of the stimulus was extracted from stimulus-present and -absent trials and compared. Both distributions of modulation amplitudes were standardized to make them approximately normal and to reduce their dependence on firing rate (Horwitz 2020). d’ was defined as the difference between the means of these two distributions, divided by their pooled standard deviation. Neurometric sensitivity, the contrast at which d’ = 1.27, was not measured.

Population d’ was defined as the d’ for an individual neuron multiplied by a population scale factor (Horwitz 2020). The population scale factor depends on the number of neurons of a given type, magnocellular or parvocellular, that are modulated by the stimulus and is defined as:

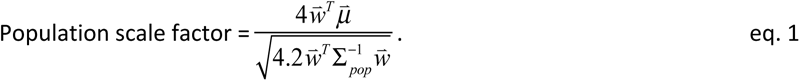

where 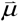 is a vector of *n* signals, with one element per neuron in the population, Σ *_pop_* is an *n* x *n* covariance matrix representing noise in the population, and 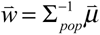 is the vector of optimal weights for population read-out. The 4 in the numerator represents the increase in signal obtained by pooling over ON and OFF mosaics in the two eyes. The 4.2 in the denominator represents the increase in noise incurred through this same pooling and includes a 0.2 that represents additional noise due to anticorrelation between ON and OFF mosaics within each eye (Ala-Laurila et al. 2011, Greschner et al. 2011, Mastronarde 1989).

To calculate 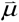 RFs were modeled as 2-dimensional Gaussian functions truncated at 2 standard deviations. A hexagonal mosaic of RFs was constructed so that each RF touched its six neighbors at the 1 standard deviation boundary (Gauthier et al. 2009). The signal carried by the *i^th^* neuron, *μ*_i_ , was defined as the integrated product of the stimulus envelope and the *i^th^* RF. The RF in the center of the hexagonal array was assumed to correspond to the neuron that was actually recorded. Σ *_pop_* , the noise covariance matrix, was constructed by assuming that every neuron was equally noisy and that correlations between neurons were equal to their RF overlap (Ala-Laurila et al. 2011, Trong & Rieke 2008). RF sizes of magnocellular neurons were taken from Derrington & Lennie (1984). Parvocellular RF sizes were taken from Watson (2014) with a 20% reduction in diameter to convert from human to macaque (Dacey & Petersen 1992).

### Behavioral SNR calculation

Threshold was defined as the contrast needed to support 82% correct choices in the contrast detection task (Gelfand & Horwitz 2018). Decisions in this task can be modeled as a comparison between draws from two independent, homoscedastic, Gaussian distributions. If the draw from the signal distribution exceeds the draw from the noise distribution, the trial is answered correctly. 82% correct is achieved when the means of the two distributions are 1.27 standard deviations apart.

Noisy estimates of detection thresholds, on which this model was based, produce noisy SNR estimates. The average cross-validated prediction error of contrast detection thresholds was 14%. A 14% change in contrast around threshold corresponds to a change from 82% correct to 79–84%, assuming a Weibull psychometric function with a slope of 3 (Wallis et al. 2013). This range corresponds to d’ values from 0.89–1.7.

### Cone current SNR calculation

The cone current model was developed by Angueyra and Rieke (2013). The implementation used in this study is identical to the one used in (Hass et al. 2015, Horwitz 2020) and is available on GitHub (https://github.com/horwitzlab/LGN-temporal-contrast-sensitivity). Each cone is modeled as a linear temporal filter, the output of which (the signal), is corrupted by additive Gaussian noise. Simulated cone currents were weighted over time and space using a filter that is identical to the signal. This resulted in two univariate, homoscedastic Gaussian distributions from which d’ was calculated (difference in means divided by standard deviation).

## Supplemental Information

### Spike train distance-based SNR analysis

The analysis of SNR presented in the main text assumes that LGN signals are at the fundamental frequency of the stimulus (F1). To examine the validity of this assumption, spike train power spectra were computed by discrete Fourier transform (Supplemental Figure 1). Parvocellular neurons, as expected, responded dominantly with an F1-modulated response component, whether the stimulus modulated the L- and M-cones in-phase (Supplemental Figure 1A) or in counterphase (Supplemental Figure 1B) (Kaplan et al. 1990). The dip in power at approximately 5 Hz is a consistent aspect of parvocellular spike trains even in the absence of contrast in the receptive field (Horwitz 2020).

Magnocellular neurons carry signatures of stimulus frequency in components of their response besides the F1. Frequency-doubled responses (F2) to high-frequency L+M stimuli were pronounced (Supplemental Figure 1C). The magnitude of the F2 component was tightly correlated with the magnitude of the F1 component across neurons within temporal frequency (mean r = 0.88, p < 0.0001, paired t-test) and across temporal frequencies within neuron (mean r = 0.81, p < 0.0001, paired t-test). The information carried by the F1 and F2 components is therefore largely redundant. A broadband increase in power at high temporal frequencies is an expected consequence of the rectangular spike counting window and the increase in spike modulation amplitude with temporal frequency (Supplemental Figure 1C & 1D)(Harris 1978).

To ask whether magnocellular spike trains carried stimulus-related signals that were missed by the analysis of F1 modulation, an auxiliary analysis was performed. Each spike train was represented as a point in a high-dimensional space, and the distance between each pair of spike trains was defined on the basis of how many spikes must be added, deleted, or moved to transform one to the other (Victor & Purpura 1997). For each stimulus condition, individual spike trains were extracted and the nine nearest neighbors identified. If five or more of these neighbors were responses to the stimulus, then the extracted spike train was classified as stimulus-present; otherwise, it was classified as stimulus-absent. These classifications were compared to ground truth to quantify correctly classified stimulus responses (hits) and incorrectly classified responses to the blank (false alarms). The signal-to-noise ratio (d’) was calculated as Φ^-1^(hit rate) - Φ^-1^(false alarm rate), where Φ^-1^ is the inverse cumulative standard normal probability density function, hit rate is the number of hits divided by the number of stimulus-present trials, and false alarm rate is the number of false alarm divided by the number of stimulus-absent trials.

The results of this analysis agreed closely with the results of the F1-based SNR analysis under most of the conditions tested (Supplemental Figure 2). The one exception was that SNR in response to high-frequency, L-M modulations in monkey 1 was considerably higher under the spike train distance-based analysis than under the F1-based analysis. In this animal, high-frequency L-M modulations often produced transient, weakly entrained responses (e.g. Figure 2B). For the comparisons made in this report, however, the assumption that signal is carried in the F1 response component is justified.

### Spike train distance-based SNR analysis methods

There are two free parameters in the calculation of SNR based on spike train distances. The first is the number of nearest neighbors used in the classification, which was set to 9. The second is the penalty associated with moving a spike 1 by second, which was set to 6 (the penalty of adding or subtracting a spike was 1). The values of these two parameters were found via a grid search that maximized decoding accuracy.

Two minor adjustments were made to d’ values calculated by this procedure to reduce variability and bias. The first adjustment avoids infinite values that would otherwise occur when a hit rate or a false alarm rate is 0 or 1. Zeros were replaced with *0.5/n* and ones were replaced with *(n-0.5)/n*, respectively, where *n* is the number trials, either stimulus-present or -absent (Stanislaw & Todorov 1999). The second correction compensates for a small downward bias caused by the fact that each trial was classified on the basis of other trials, which are likely to be of the opposite type. Consider an urn containing equal numbers of red and black balls: the nearest neighbors of a red ball, not including itself, are more likely to be black than red. To correct for this fact, Φ^-1^ (E(hit rate)) - Φ^-1^ (E(false alarm rate)) was subtracted from d’, where E(hit rate) and E(false alarm rate) are the expected values of the hit rate and false alarm rate under the ball and urn model.

## Supplemental Figure legends

**Supplemental Figure 1.** Spectral analysis of LGN spike trains. Power spectra are shown for parvocellular responses to L+M stimuli (**A**), parvocellular responses to L-M stimuli (**B**), magnocellular responses to L+M stimuli (**C**), and magnocellular responses to L-M stimuli (**D**). The fundamental frequency of each stimulus is shown along the abscissa (triangles). Frequency-doubled responses in (C) are indicated by black arrows.

**Supplemental Figure 2.** Magnocellular signal-to-noise ratio (d’) in response to L+M modulations (black) and L-M modulations (magenta). d’ was computed from responses at the fundamental frequency of the stimulus (closed symbols) and from the performance of a k-nearest neighbors spike train classifier (open symbols). Symbols represent means across neurons, and shaded bands represent ± 1 standard error of the mean. Data are from monkey 1 (**A**) and monkey 2 (**B**).

**Supplemental Figure 3.** Analysis of spike counting window on population signal-to-noise ratio. Population signal-to-noise ratio was calculated from parvocellular responses to L-M modulations (magenta) and magnocellular responses to L+M modulations (black). Spikes were counted for 0.05, 0.1, 0.2, 0.4, or 0.66 s (columns), starting 0, 0.05, 0.1, 0.15, 0.2, 0.25, or 0.3 s after stimulus onset (rows). Symbols represent means across neurons, and shaded bands represent ±1 standard error of the mean. Horizontal lines indicate signal-to-noise ratios of 0 (gray) and 1.27 (dashed).

**Supplemental Figure 4.** Signal-to-noise ratio of individual LGN neurons and the cones assumed to be inside their receptive fields. Plotting conventions are as in Figure 5, but population scaling has not been performed on the individual neuronal d’ values, and only the cones inside the receptive field of each recorded LGN neuron were simulated.

## References

Ala-Laurila P, Greschner M, Chichilnisky EJ, Rieke F. 2011. Cone photoreceptor contributions to noise and correlations in the retinal output. Nat Neurosci 14: 1309–16

Alitto HJ, Usrey WM. 2008. Origin and dynamics of extraclassical suppression in the lateral geniculate nucleus of the macaque monkey. Neuron 57: 135–46

Anderson SJ, Mullen KT, Hess RF. 1991. Human peripheral spatial resolution for achromatic and chromatic stimuli: limits imposed by optical and retinal factors. The Journal of physiology 442: 47–64

Angueyra JM, Rieke F. 2013. Origin and effect of phototransduction noise in primate cone photoreceptors. Nat Neurosci 16: 1692–700

Banks MS, Geisler WS, Bennett PJ. 1987. The physical limits of grating visibility. Vision research 27: 1915–24

Banks MS, Sekuler AB, Anderson SJ. 1991. Peripheral spatial vision: limits imposed by optics, photoreceptors, and receptor pooling. *Journal of the Optical Society of America A*, Optics and image science 8: 1775–87

Barlow HB. 1957. Increment thresholds at low intensities considered as signal/noise discriminations. The Journal of physiology 136: 469–88

Baudin J, Angueyra JM, Sinha R, Rieke F. 2019. S-cone photoreceptors in the primate retina are functionally distinct from L and M cones. Elife 8

Benardete EA, Kaplan E. 1997a. The receptive field of the primate P retinal ganglion cell, I: Linear dynamics. Vis Neurosci 14: 169–85

Benardete EA, Kaplan E. 1997b. The receptive field of the primate P retinal ganglion cell, II: Nonlinear dynamics. Vis Neurosci 14: 187–205

Benardete EA, Kaplan E. 1999a. The dynamics of primate M retinal ganglion cells. Vis Neurosci 16: 355–68

Benardete EA, Kaplan E. 1999b. Dynamics of primate P retinal ganglion cells: responses to chromatic and achromatic stimuli. J Physiol 519 Pt 3: 775–90

Brainard DH. 1997. The Psychophysics Toolbox. Spat Vis 10: 433–6

Carvalho LS, Pessoa DMA, Mountford JK, Davies WIL, Hunt DM. 2017. The Genetic and Evolutionary Drives behind Primate Color Vision. Frontiers in Ecology and Evolution 5

Chaparro A, Stromeyer CF, Huang EP, Kronauer RE, Eskew RT. 1993. Colour Is What the Eye Sees Best. Nature 361: 348–50

Conway BR, Livingstone MS. 2006. Spatial and temporal properties of cone signals in alert macaque primary visual cortex. J Neurosci 26: 10826–46

Cottaris NP, Wandell BA, Rieke F, Brainard DH. 2020. A computational observer model of spatial contrast sensitivity: Effects of photocurrent encoding, fixational eye movements, and inference engine. Journal of vision 20: 17

Croner LJ, Kaplan E. 1995. Receptive fields of P and M ganglion cells across the primate retina. Vision Res 35: 7–24

Curcio CA, Sloan KR, Jr., Packer O, Hendrickson AE, Kalina RE. 1987. Distribution of cones in human and monkey retina: individual variability and radial asymmetry. Science 236: 579–82

Dacey DM. 1993. The mosaic of midget ganglion cells in the human retina. The Journal of neuroscience : the official journal of the Society for Neuroscience 13: 5334–55

Dacey DM, Petersen MR. 1992. Dendritic field size and morphology of midget and parasol ganglion cells of the human retina. Proc Natl Acad Sci U S A 89: 9666–70

De A, Horwitz GD. 2021. Coding of chromatic spatial contrast by macaque V1 neurons. bioRxiv

De Lange Dzn H. 1958. Research into the dynamic nature of the human fovea→ cortex systems with intermittent and modulated light. I. Attenuation characteristics with white and colored light. Journal of the Optical Society of America 48: 777–84

Derrington AM, Lennie P. 1984. Spatial and temporal contrast sensitivities of neurones in lateral geniculate nucleus of macaque. J Physiol 357: 219–40

Eickhoff SB, Rottschy C, Zilles K. 2007. Laminar distribution and co-distribution of neurotransmitter receptors in early human visual cortex. Brain structure & function 212: 255–67

Engel S, Zhang X, Wandell B. 1997. Colour tuning in human visual cortex measured with functional magnetic resonance imaging. Nature 388: 68–71

Enroth-Cugell C, Robson JG, Schweitzer-Tong DE, Watson AB. 1983. Spatio-temporal interactions in cat retinal ganglion cells showing linear spatial summation. The Journal of physiology 341: 279–307

Gauthier JL, Field GD, Sher A, Shlens J, Greschner M, et al. 2009. Uniform signal redundancy of parasol and midget ganglion cells in primate retina. The Journal of neuroscience : the official journal of the Society for Neuroscience 29: 4675–80

Geisler WS. 1989. Sequential Ideal-Observer Analysis of Visual Discriminations. Psychological Review 96: 267–314

Geisler WS. 2011. Contributions of ideal observer theory to vision research. Vision research 51: 771–81

Gelfand EC, Horwitz GD. 2018. Model of parafoveal chromatic and luminance temporal contrast sensitivity of humans and monkeys. J Vis 18: 1

Green DM, Swets JA. 1966. Signal detection theory and psychophysics. New York: Wiley

Greschner M, Shlens J, Bakolitsa C, Field GD, Gauthier JL, et al. 2011. Correlated firing among major ganglion cell types in primate retina. J Physiol 589: 75–86

Gur M, Snodderly DM. 1997. A dissociation between brain activity and perception: chromatically opponent cortical neurons signal chromatic flicker that is not perceived. Vision Res 37: 377–82

Harris FJ. 1978. On the use of windows for harmonic analysis with the discrete Fourier transform Proceedings of the IEEE 66: 51–83

Hass CA, Angueyra JM, Lindbloom-Brown Z, Rieke F, Horwitz GD. 2015. Chromatic detection from cone photoreceptors to V1 neurons to behavior in rhesus monkeys. J Vis 15: 1

Hawken MJ, Shapley RM, Grosof DH. 1996. Temporal-frequency selectivity in monkey visual cortex. Vis Neurosci 13: 477–92

Horwitz GD. 2020. Temporal information loss in the macaque early visual system. Plos Biology 18

Jiang Y, Purushothaman G, Casagrande VA. 2015a. A computational relationship between thalamic sensory neural responses and contrast perception. Front Neural Circuits 9: 54

Jiang Y, Yampolsky D, Purushothaman G, Casagrande VA. 2015b. Perceptual decision related activity in the lateral geniculate nucleus. J Neurophysiol 114: 717–35

Jiang Y, Zhou K, He S. 2007. Human visual cortex responds to invisible chromatic flicker. Nat Neurosci 10: 657–62

Kaplan E, Benardete E. 2001. The dynamics of primate retinal ganglion cells. Prog Brain Res 134: 17–34

Kaplan E, Lee BB, Shapley R. 1990. New views of primate retinal function. Progress in retinal research 9: 273–336

Kaplan E, Purpura K, Shapley RM. 1987. Contrast affects the transmission of visual information through the mammalian lateral geniculate nucleus. J Physiol 391: 267–88

Kaplan E, Shapley RM. 1986. The primate retina contains two types of ganglion cells, with high and low contrast sensitivity. Proc Natl Acad Sci U S A 83: 2755–7

Kelly DH, van Norren D. 1977. Two-band model of heterochromatic flicker. J Opt Soc Am 67: 1081–91

King-Smith PE, Carden D. 1976. Luminance and opponent-color contributions to visual detection and adaptation and to temporal and spatial integration. J Opt Soc Am 66: 709–17

Kleiner M, Brainard DH, Pelli DG. 2007. What’s new in Psychtoolbox-3. Presented at Perception 36 ECVP Abstract Supplement

Koenig D, Hofer H. 2011. The absolute threshold of cone vision. Journal of Vision 11

Kremkow J, Alonso JM. 2018. Thalamocortical Circuits and Functional Architecture. Annu Rev Vis Sci 4: 263–85

Krolak-Salmon P, Henaff MA, Tallon-Baudry C, Yvert B, Guenot M, et al. 2003. Human lateral geniculate nucleus and visual cortex respond to screen flicker. Ann Neurol 53: 73–80

Krukowski AE, Miller KD. 2001. Thalamocortical NMDA conductances and intracortical inhibition can explain cortical temporal tuning. Nat Neurosci 4: 424–30

Krukowski AE, Troyer TW, Miller KD. 2001. A model of visual cortical temporal frequency tuning. Neurocomputing 38: 1379–83

Lankheet MJ, Lennie P, Krauskopf J. 1998. Temporal-chromatic interactions in LGN P-cells. Vis Neurosci 15: 47–54

Lee BB, Pokorny J, Smith VC, Martin PR, Valberg A. 1990. Luminance and chromatic modulation sensitivity of macaque ganglion cells and human observers. J Opt Soc Am A 7: 2223–36

Lee BB, Sun H. 2009. The chromatic input to cells of the magnocellular pathway of primates. J Vis 9: 15 1–18

Legall D. 1991. Mpeg - a Video Compression Standard for Multimedia Applications. Communications of the Acm 34: 46–58

Lester RA, Clements JD, Westbrook GL, Jahr CE. 1990. Channel kinetics determine the time course of NMDA receptor-mediated synaptic currents. Nature 346: 565–7

Lindbloom-Brown Z, Tait LJ, Horwitz GD. 2014. Spectral sensitivity differences between rhesus monkeys and humans: Implications for neurophysiology. Journal of Neurophysiology

Martin PR, Lee BB, White AJ, Solomon SG, Ruttiger L. 2001. Chromatic sensitivity of ganglion cells in the peripheral primate retina. Nature 410: 933–6

Mastronarde DN. 1989. Correlated firing of retinal ganglion cells. Trends Neurosci 12: 75–80

Merigan WH. 1980. Temporal modulation sensitivity of macaque monkeys. Vision Res 20: 953–9

Merigan WH, Eskin TA. 1986. Spatio-temporal vision of macaques with severe loss of P beta retinal ganglion cells. Vision Res 26: 1751–61

Merigan WH, Maunsell JH. 1990. Macaque vision after magnocellular lateral geniculate lesions. Vis Neurosci 5: 347–52

Mullen KT, Kingdom FAA. 2002. Differential distributions of red-green and blue-yellow cone opponency across the visual field. Visual neuroscience 19: 109–18

Mullen KT, Sakurai M, Chu W. 2005. Does L/M cone opponency disappear in human periphery? Perception 34: 951–9

Owsley C. 2011. Aging and vision. Vision Res 51: 1610–22

Packer O, Hendrickson AE, Curcio CA. 1989. Photoreceptor topography of the retina in the adult pigtail macaque (Macaca nemestrina). J Comp Neurol 288: 165–83

Pelli DG. 1997. The VideoToolbox software for visual psychophysics: transforming numbers into movies. Spat Vis 10: 437–42

Pokorny J, Smith VC, Lee BB, Yeh T. 2001. Temporal sensitivity of macaque ganglion cells to lights of different chromaticity. Color Research and Application 26: S140–S44

Purpura K, Tranchina D, Kaplan E, Shapley RM. 1990. Light adaptation in the primate retina: analysis of changes in gain and dynamics of monkey retinal ganglion cells. Vis Neurosci 4: 75–93

Rider AT, Henning GB, Stockman A. 2019. Light adaptation controls visual sensitivity by adjusting the speed and gain of the response to light. PloS one 14: e0220358

Robson JG. 1966. Spatial and Temporal Contrast-Sensitivity Functions of Visual System. Journal of the Optical Society of America 56: 1141-&

Rosenthal I, Ratnasingam S, Haile T, Eastman S, Fuller-Deets J, Conway BR. 2018. Color statistics of objects, and color tuning of object cortex in macaque monkey. Journal of vision 18: 1

Rovamo J, Kukkonen H, Raninen A, Donner K. 2003. Efficiency of temporal integration of sinusoidal flicker. Invest Ophthalmol Vis Sci 44: 5049–55

Rovamo J, Raninen A, Lukkarinen S, Donner K. 1996. Flicker sensitivity as a function of spectral density of external white temporal noise. Vision Res 36: 3767–74

Ruderman DL, Cronin TW, Chiao CC. 1998. Statistics of cone responses to natural images: implications for visual coding. Journal of the Optical Society of America a-Optics Image Science and Vision 15: 2036–45

Schiller PH, Logothetis NK, Charles ER. 1990. Role of the color-opponent and broad-band channels in vision. Vis Neurosci 5: 321–46

Shady S, MacLeod DIA, Fisher HS. 2004. Adaptation from invisible flicker. Proceedings of the National Academy of Sciences of the United States of America 101: 5170–3

Sinha R, Hoon M, Baudin J, Okawa H, Wong ROL, Rieke F. 2017. Cellular and Circuit Mechanisms Shaping the Perceptual Properties of the Primate Fovea. Cell 168: 413–26 e12

Smith VC, Bowen RW, Pokorny J. 1984. Threshold temporal integration of chromatic stimuli. Vision Res 24: 653–60

Snowden RJ, Hess RF. 1992. Temporal frequency filters in the human peripheral visual field. Vision Res 32: 61–72

Snowden RJ, Hess RF, Waugh SJ. 1995. The processing of temporal modulation at different levels of retinal illuminance. Vision Res 35: 775–89

Solomon SG, Lee BB, White AJR, Ruttiger L, Martin PR. 2005. Chromatic organization of ganglion cell receptive fields in the peripheral retina. The Journal of neuroscience : the official journal of the Society for Neuroscience 25: 4527–39

Solomon SG, Martin PR, White AJ, Ruttiger L, Lee BB. 2002. Modulation sensitivity of ganglion cells in peripheral retina of macaque. Vision Res 42: 2893–8

Sperling HG, Crawford ML, Espinoza S. 1978. Threshold spectral sensitivity of single neurons in the lateral geniculate nucleus and of performing monkeys. Mod Probl Ophthalmol 19: 2–18

Stanislaw H, Todorov N. 1999. Calculation of signal detection theory measures. Behav Res Methods Instrum Comput 31: 137–49

Stavros KA, Kiorpes L. 2008. Behavioral measurement of temporal contrast sensitivity development in macaque monkeys (Macaca nemestrina). Vision Res 48: 1335–44

Stockman A, Henning GB, Anwar S, Starba R, Rider AT. 2018. Delayed cone-opponent signals in the luminance pathway. J Vis 18: 6

Stockman A, Langendorfer M, Smithson HE, Sharpe LT. 2006. Human cone light adaptation: from behavioral measurements to molecular mechanisms. J Vis 6: 1194–213

Swanson WH, Ueno T, Smith VC, Pokorny J. 1987. Temporal modulation sensitivity and pulse- detection thresholds for chromatic and luminance perturbations. J Opt Soc Am A 4: 1992–2005

Tolhurst DJ, Dean AF. 1990. The effects of contrast on the linearity of spatial summation of simple cells in the cat’s striate cortex. Exp Brain Res 79: 582–8

Trong PK, Rieke F. 2008. Origin of correlated activity between parasol retinal ganglion cells. Nat Neurosci 11: 1343–51

Troyer TW, Krukowski AE, Priebe NJ, Miller KD. 1998. Contrast-invariant orientation tuning in cat visual cortex: thalamocortical input tuning and correlation-based intracortical connectivity. J Neurosci 18: 5908–27

Tyler CW. 1981. Specific deficits of flicker sensitivity in glaucoma and ocular hypertension. Invest Ophthalmol Vis Sci 20: 204–12

Van der Horst GJC. 1969. Chromatic Flicker. Journal of the Optical Society of America 59: 1213–17

Victor JD, Purpura KP. 1997. Metric-space analysis of spike trains: Theory, algorithms and application. Network-Computation in Neural Systems 8: 127–64

Vul E, MacLeod DI. 2006. Contingent aftereffects distinguish conscious and preconscious color processing. Nat Neurosci 9: 873–4

Wallis SA, Baker DH, Meese TS, Georgeson MA. 2013. The slope of the psychometric function and non-stationarity of thresholds in spatiotemporal contrast vision. Vision Res 76: 1–10

Watson AB. 2014. A formula for human retinal ganglion cell receptive field density as a function of visual field location. J Vis 14

Wiesel TN, Hubel DH. 1966. Spatial and chromatic interactions in the lateral geniculate body of the rhesus monkey. Journal of Neurophysiology 29: 1115–56

Williams PE, Mechler F, Gordon J, Shapley R, Hawken MJ. 2004. Entrainment to video displays in primary visual cortex of macaque and humans. J Neurosci 24: 8278–88

Zhigalov A, Herring JD, Herpers J, Bergmann TO, Jensen O. 2019. Probing cortical excitability using rapid frequency tagging. Neuroimage 195: 59–66

