## Supplementary figures and images for "Temporal filtering of luminance and chromaticity in macaque visual cortex"

### Supplemental Figure 1

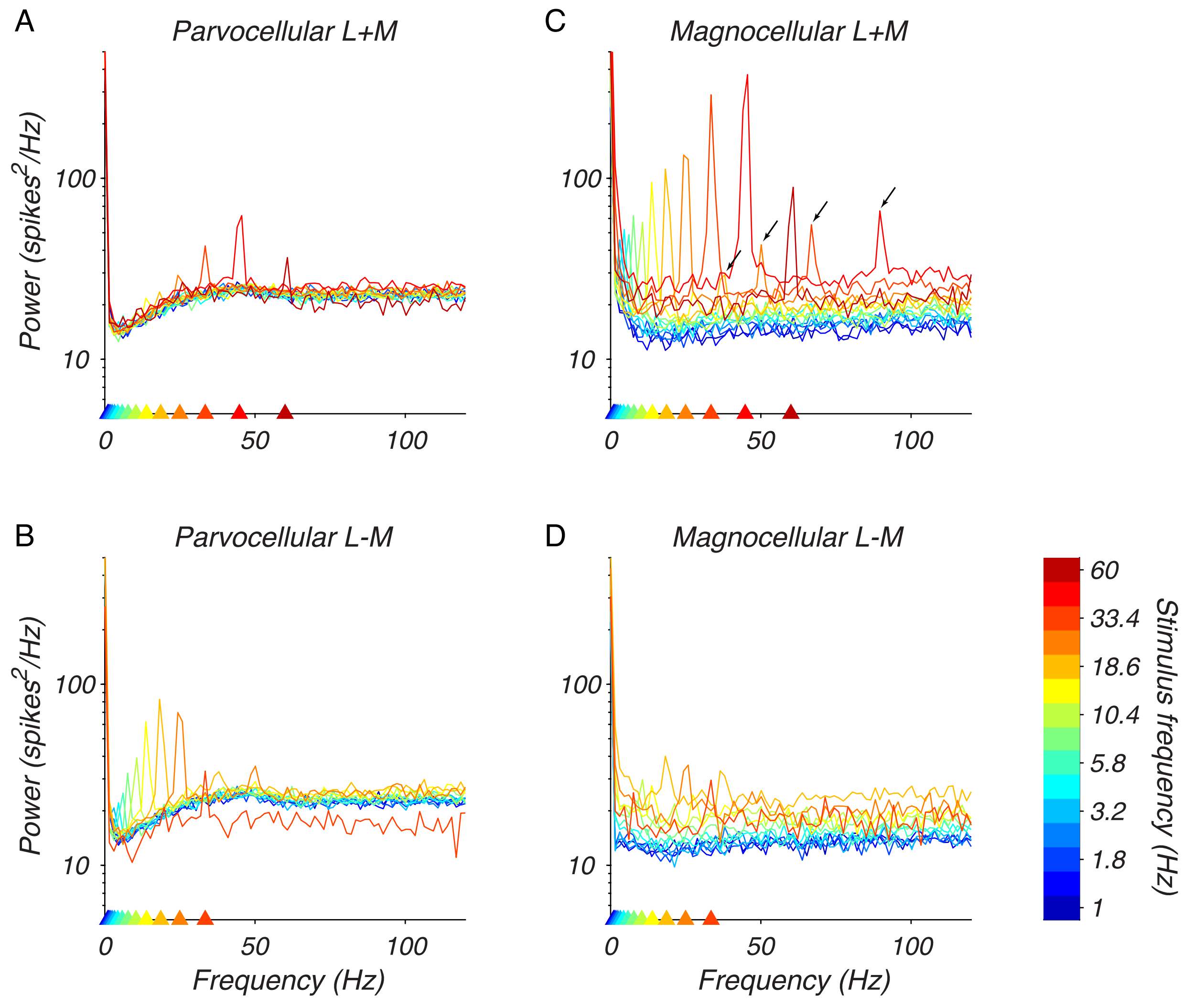

### Supplemental Figure 2

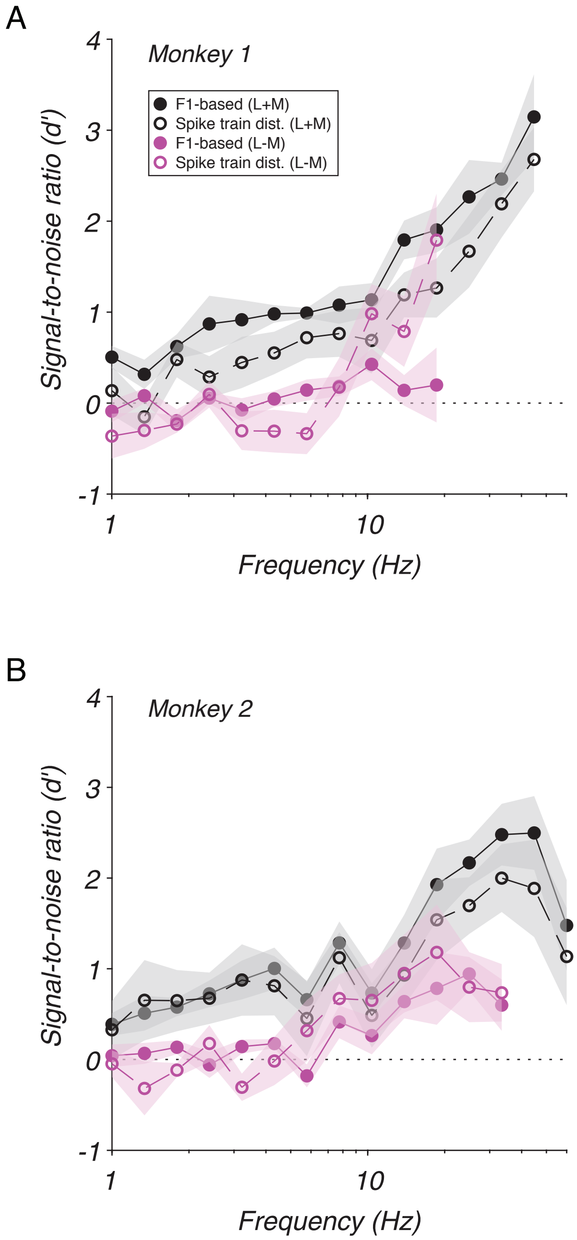

### Supplemental Figure 3

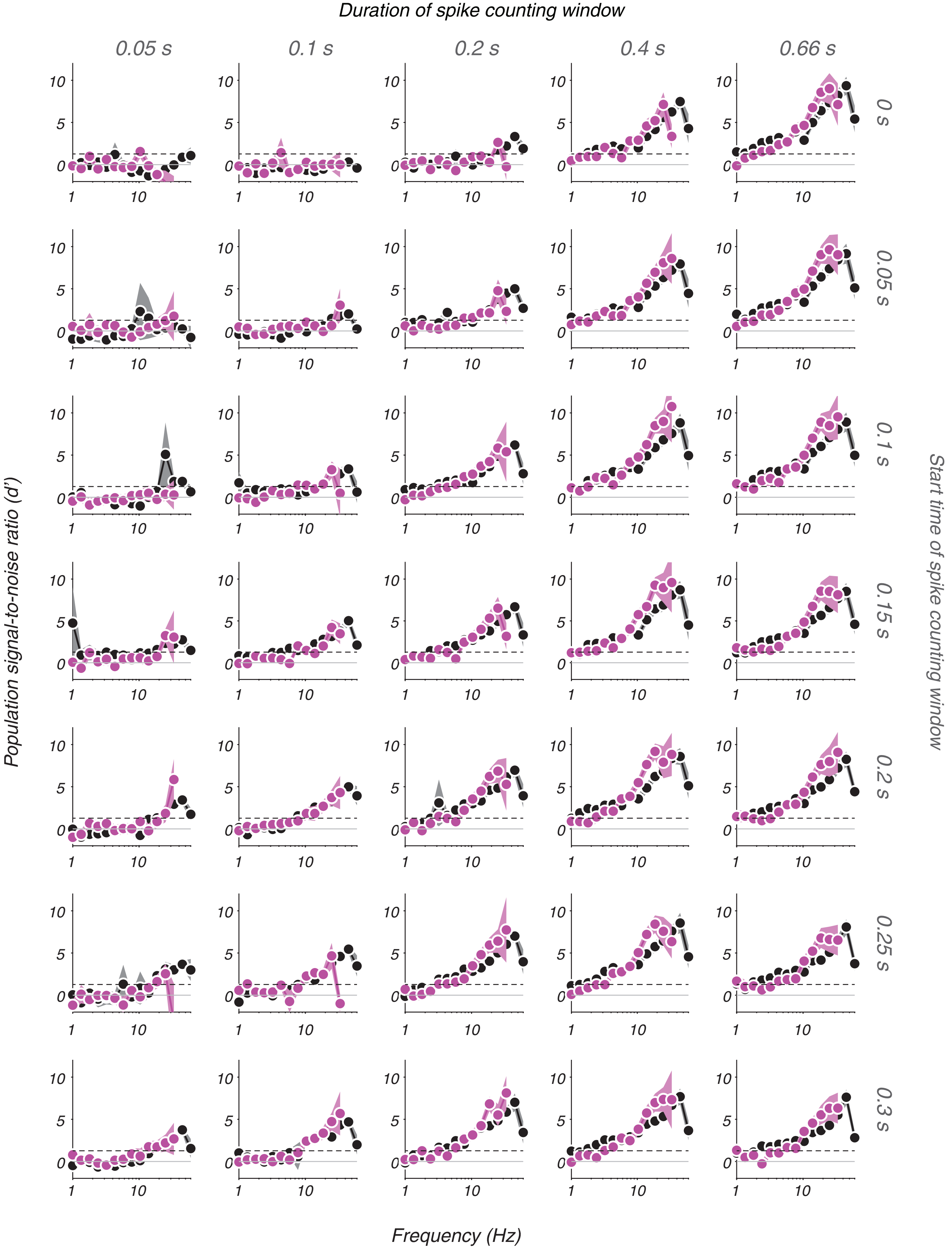

### Supplemental Figure 4

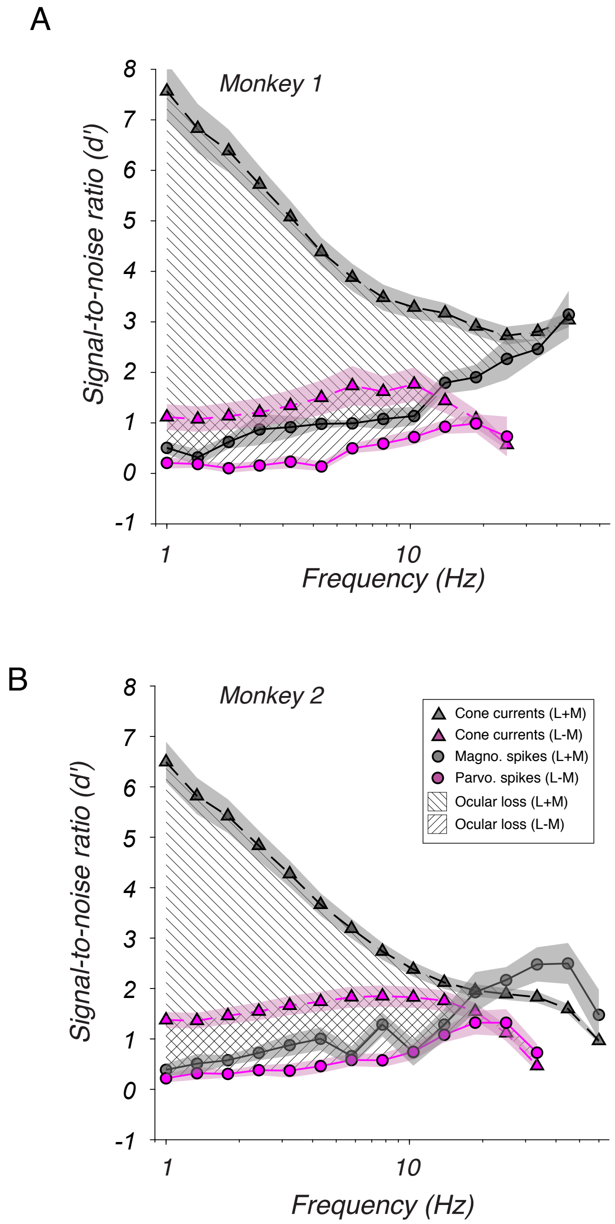
